# A mitotic bookmark coordinates transcription and replication

**DOI:** 10.1101/2025.06.29.662044

**Authors:** Chun-Yi Cho, Patrick H. O’Farrell

**Affiliations:** Department of Biochemistry and Biophysics, University of California, San Francisco, San Francisco, CA 94158, USA

## Abstract

Collisions between advancing replication forks and elongating transcripts pose a universal threat. During the rapid nuclear division cycles in early *Drosophila* embryos, coordinating transcription and replication is critical to reduce the risk of collisions. In each cycle, replication begins immediately after mitosis, while transcription starts 3 minutes later, overlapping with replication for the remainder of interphase. We previously showed that transcription depends on the coactivator Brd4, which forms hubs at active genes. Here, we show that Brd4 persists on mitotic chromosomes as bookmarks of transcriptional activity and, upon anaphase entry, recruits the replication activator Cdc7 to specify early-replicating genomic regions in the following interphase. Additionally, Cdc7 activity removes Brd4 bookmarks such that post-mitotic transcription occurs only after a new round of Brd4 hub assembly. Early initiation of replication while deferring initiation of transcription is proposed to allow unimpeded transcriptional elongation behind advancing replication forks. Supporting this, inhibiting Cdc7 delayed replication, stabilized Brd4 bookmarks, and resulted in premature transcription with elongation defects. We propose that Cdc7 triggers a functional switch in Brd4 that enforces temporal ordering of the initiation of transcription and replication, thereby minimizing collisions. This switching process might underlie the widespread correlation between transcriptional activity and early replication.

## INTRODUCTION

All dividing cells face the risk of collisions between transcription and replication machineries (1, 2). Such collisions disrupt both processes, leading to DNA damage and genome instability (3–5). Despite this risk, the central dogma has been conserved throughout evolution, with DNA serving as the template for both transcription and replication. This suggests the existence of mechanisms that prevent or at least minimize these collisions. Indeed, studies in diverse organisms—including ciliates, yeast, and humans—revealed that transcription and replication normally do not occur simultaneously at the same genomic region (6–8). However, the mechanisms underlying this coordination remain largely undefined.

Early embryogenesis in *Drosophila* provides an excellent model for exploring this question. After fertilization, the embryo undergoes 13 synchronous cell cycles as a syncytium (<u>n</u>uclear division <u>c</u>ycles, NCs), rapidly alternating between S and M phases without intervening gap phases or cytokinesis. An early wave of zygotic transcription begins in NC8 and continues through NC13 (9), with transcripts initiating about 3 minutes into each S phase and aborting upon entry into mitosis (10). Despite overlapping with intense replication activity, genes are successfully transcribed during these short S phases. We recently found that, during NC12, transcription depends on replication (11). This coupling of transcription to replication was suggested to contribute to a “replication-first” strategy, in which transcription initiates only on replicated DNA segments and can follow replication forks unimpeded.

Developmental changes in the cell cycle complicate a replication-first strategy. In early *Drosophila* embryos, the S-phase duration progressively lengthens, with a roughly two-fold increase from NC12 (8-9 minutes) to NC13 (15-16 minutes) and a dramatic increase to about an hour in NC14 (12, 13). This lengthening is largely driven by the emergence of a replication-timing program, where different genomic regions initiate replication at distinct times during S phase, in contrast to the near-synchronous initiation throughout the genome in the earliest short cycles (12, 14). If the coupling of transcription to replication meant that transcription had to await local replication of its template during a prolonged S phase, the onset of transcription would be significantly delayed throughout much of the genome, unless transcribed loci are preferentially replicated early. Given that transcription still initiates about 3 minutes into NC13 and considering that the transcribed genes generally replicate early in S phase (14), we hypothesized the existence of a mechanism that promotes early replication of transcribed loci in the syncytial embryo.

The chromatin environment regulates DNA-templated processes. One key mechanism involves the recognition of histone marks by specialized reader proteins (15). Among these, bromodomain-containing proteins are the primary readers of histone acetylation, a hallmark of open chromatin and active genes (16, 17). Members of the bromodomain and extraterminal domain (BET) family, such as Brd4, have been extensively studied for their roles in both transcription and replication. Brd4 promotes transcription by recruiting the positive transcription elongation factor b (P-TEFb)(18, 19), while also advancing replication timing by recruiting the replication initiation factor Treslin (20). Notably, Brd4 is one of the few transcriptional regulators that remain bound to select loci during mitosis. Localized mitotic binding is thought to serve as a bookmark, preserving a memory of gene activity from the previous cell cycle (21–25). Mitotic bookmarks are generally associated with the rapid reactivation of gene expression in daughter cells (26), but whether their function extends to other biological processes is unknown.

Recent studies show that the *Drosophila* homologs of Brd4/Fs(1)h and its upstream regulator, the lysine acetyltransferase CBP/Nejire, activate early zygotic transcription (27–29). These factors act downstream of pioneer transcription factors, including Zelda for non-histone genes and an unidentified factor for the replication-dependent histone genes (27). Brd4 forms dynamic clusters that emerge and disperse during progression from NC12 to NC13 (27): clusters appear about 2 minutes into S phase 12, persist through mitosis, disperse rapidly at the onset of the S phase 13, and then reappear after another 2-minute delay. When they first emerge in NC12, Brd4 clusters function as “transcriptional hubs” to enrich RNA polymerase II and fuel a burst of transcription about 3 min into S phase. On mitotic chromosomes, Brd4 clusters persist prominently at the histone gene repeats, as well as at other smaller foci that are Zelda-dependent and likely correspond to early expressed non-histone genes (27). While mitotic binding is consistent with a role in bookmarking active genes, these clusters disperse rapidly after mitosis and must reassemble before transcription begins 3 minutes later, suggesting that any contribution of mitotically stable Brd4 to transcriptional reactivation in a subsequent NC is indirect. Given the absence of gap phases and the immediate onset of DNA replication after mitosis, we hypothesized that mitotic Brd4 clusters serve to promote early replication timing in NC13 and NC14. Advancing replication timing can in principle sustain the replication-first order while providing bookmarked loci the opportunity to initiate transcription early despite overall slowing of S phase.

Here, we uncover a role of mitotic bookmarks in promoting early replication of specific genomic regions in syncytial *Drosophila* embryos. We demonstrate that Brd4 hubs undergo a functional switch from a role in transcription when they first emerge in interphase to one that promotes replication at the onset of the next S phase. Using live imaging, we show that Brd4 clusters persist on mitotic chromosomes and recruit the replication initiation factor, Cdc7 kinase, to select loci during entry into S phase. This mechanism links transcriptional activation in one cycle to replication timing in the next cycle, ensuring that active genes are replicated early in a prolonged S phase. Importantly, the early recruitment of Cdc7 triggered the dispersal of Brd4 bookmarks, thereby enforcing a dependency of transcription on a new round of transcriptional hub assembly and ensuring that transcription follows replication. Inhibiting Cdc7 stabilized Brd4 bookmarks and delayed replication, changes that were associated with precocious RNAPII recruitment and disrupted transcriptional elongation. We suggest that the ordering of events enforced by Cdc7 function delays transcription such that it begins only after replication forks have cleared the region, and that the disruption of this coordination leads to collisions interfering with transcript elongation. Finally, we propose that chromatin bookmarks have a switchable, dual function that provides an explanation for the correlation between early replication and transcriptional activity in eukaryotes.

## MATERIALS AND METHODS

### Fly stocks

*Drosophila melanogaster* were maintained on standard cornmeal-yeast medium at 25°C. Flies were transferred to egg-laying cages 2-3 days before embryo collection for experiments. Fly lines used in this study are listed in Supplementary Table 1.

### Embryo mounting for live imaging

Embryos were collected from females in egg-laying cage onto a grape juice agar plate with yeast paste, dechorionated with 40% bleach in a basket, washed with water, and then transferred to a new grape juice agar plate. Embryos were aligned, glued to a coverslip, and then covered with halocarbon oil (1:1 mixture of halocarbon oil 27 and 700) for microscopy.

### Spinning disk confocal microscopy

Imaging was performed on an Olympus IX70 microscope equipped with PerkinElmer Ultraview Vox confocal system. Movies of the *hunchback* transcriptional reporters were acquired using a 60x/1.40 oil objective with the pixel binning set to 1×1, while all other movies were acquired using a 100x/1.40 oil objective with the pixel binning set to 2×2. Data were acquired using Volocity 6 software (Quorum Technologies). Z-stacks were recorded with a 0.5 μm step size at each time point. Fluorophores were excited with 488 and 561 nm laser lines. Appropriate emission filters were used except when performing two-color imaging of mKate2-Brd4 and Cdc7-EGFP, which exhibited low fluorescent intensity and required fast frame rate. For these cases, the “fast sequential” mode was used to skip emission filters, and care was taken to make sure that there was negligible bleed. Images in the same set of experiments were acquired using the same configuration, and laser power was calibrated using a laser power meter (Thorlabs) before each imaging session.

### Microinjection

Once glued to the coverslip, embryos were slightly dehydrated in a desiccation chamber for 7-9 minutes before being covered in halocarbon oil and subjected to microinjection. JQ1 (abcam, #146612) supplied as 10 mM solution in DMSO was diluted fresh with water to 0.1 mM for injection. α-amanitin (Sigma-Aldrich, #A2263) was dissolved in water and injected at 0.5 mg/ml. XL413 (Sigma-Aldrich, #SML1401) was dissolved in water at 5 mg/mL as stock solution and diluted fresh to 0.5 mg/ml for injection. ECFP-Geminin was injected at 20 mg/ml in 40 mM HEPES pH 7.4, 150 mM KCl buffer. To inject XL413 precisely during mitosis, live confocal microscopy was used for staging. When the embryo entered mitosis, the entire slide was removed from the microscope for injection on an adjacent station and returned for imaging within 2-3 minutes.

### Pulse labeling of Cy5-dUTP

Embryos injected with 0.1 mM Cy5-dUTP (Sigma-Aldrich, #GEPA55022) were incubated for 3 minutes and then recovered for fixation as described (11). Briefly, embryos were washed off the glue by heptane, fixed by a 1:1 mixture of heptane and 37% formaldehyde, and dehydrated by methanol. Embryos were transferred with glass pipette onto a nylon mesh, washed with PBS, and transferred onto a strip of double-sided tape in a glass dish. A dissecting probe was used to gently rolled the embryos on the tape to remove the vitelline membrane, and then PBST (0.3% tween 20) was used to transfer the embryos to a 0.5-mL tube. Embryos were further washed with PBST for 15 minutes twice and mounted for imaging.

### Fly crosses for germline RNAi experiments

To generate females for embryo collection in the *CBP* RNAi experiments, we first set up the following P0 cross: *mKate2-Brd4, Cdc7-EGFP; UASp-shRNA* (females) x *mKate2-Brd4, Cdc7-EGFP; Mat-tub-Gal4 (males)*. The resulting F1 progeny expressed the shRNA targeting *white* (control) or *CBP* in the female germline under the control of *Mat-tub-Gal4*. The progeny were crossed with their siblings and transferred to egg-laying cages for embryo collection. To increase the Gal4/UASp induction of shRNA expression in the egg laying females, F1 progeny were grown at 27°C from larval stage onward.

### Fly crosses for imaging of *hunchback* transcriptional reporters

The following crosses were set up to collect embryos for imaging of transcriptional reporters.

1. *EGFP-Rpb3; nos-MCP-mCherry* (females) x *hbP2-MS2-lacZ* (males)
2. *nos-NLS-mCherry-PCP, His2Av-eBFP2/+ (II); MCP-GFP/+ (III)* (females) x *hbP2-MS2-lacZ-PP7*

Flies were crossed on standard medium for 2-3 days before transferring to egg-laying cages. Females for Cross #2 were collected from a separate cross between wild type and *nos-NLS-mCherry-PCP, His2Av-eBFP2/CyO (II); MCP-GFP (III)*.

### Molecular cloning and CRISPR-Cas9 genome editing

To knock-in mKate2 at the N-terminus of endogenous Brd4 (*Fs*(*1*)*h*), the mKate2 DNA fragment (optimized with *Drosophila* codons) was synthesized by Integrated DNA Technologies (IA) and used to replace the *sfGFP* sequence in the donor plasmid for *sfGFP-Brd4* by Gibson assembly. The sgRNA-expressing and donor plasmids were sent to Rainbow Transgenic Flies (Camarillo, CA) for microinjection. After injection, surviving adults were crossed to *yw, N/FM7c* balancer flies and screened by PCR for successful knock-in. Transformants were backcrossed with wild-type flies at least three times before performing experiments. The knock-in of *mNeonGreen* at the N-terminus of endogenous *CBP* (*neijire*) was designed and performed by WellGenetics (Taiwan). Briefly, the mNeonGreen-loxP-3xP3-RFP-loxP cassette was first inserted by CRISPR-Cas9 using sgRNA targeting the 5’ end of *CBP* (5’-AGACGAACCGCCCCAAAAGC-3’), followed by Cre-mediated excision through genetic crossing. A linker peptide (RSITSYNVCYTKLSAS) was left between mNeonGreen and CBP. **Image processing and presentation**

Data acquired in Volocity were exported as TIFF files and further processed using FIJI/ImageJ. Z-stack images were first converted to maximal intensity projections, followed by background subtraction using a rolling-ball radius of 50 pixels. Regions containing nuclei located within the Z-stack (excluding peripheral ones) were cropped for downstream analysis. Figures show data of one representative experiment from at least three independent replicates.

### Image analysis

Analysis was performed using custom FIJI macros and Python scripts, using standard and open-sourced libraries including SciPy, NumPy, scikit-image, Trackpy, and napari (30–34).

### Quantification of mKate2-Brd4 and Cdc7-EGFP movies

Binary masks for mitotic chromosomes and interphase nuclei were generated by summing the fluorescence intensities from both channels followed by Gaussian blurring. To reduce the impact of bright foci on segmentation, intensities above a manually determined threshold were replaced with a lower constant value. Segmentation was then achieved using Otsu’s thresholding method. For comparing the dynamics between nuclear cycles in untreated embryos, the mean and variance of mKate2-Brd4 and Cdc7-EGFP fluorescence intensities within the combined mask area were computed at each time point. To compare dynamics between treatments, fluorescence intensities were measured within individual nuclei, and data were pooled from multiple embryos for analysis.

### Quantification of EGFP-Rpb3 and MCP-mCherry movies

Binary masks for the nuclei were generated using EGFP-Rpb3 signals. The especially bright signals at HLBs were reduced by replacing intensities above a specified threshold with a lower constant value, followed by Gaussian blurring to smooth the images. Initial segmentation was achieved using Otsu’s thresholding method, and the nuclear masks were dilated to capture MCP foci at the periphery. A watershed transformation was applied to separate touching nuclei, and a final filtering step removed small nuclei (touching the top or bottom of Z-stacks) and those touching X-Y borders. Nuclei were tracked using the tracky package, and only the nuclei detected across all frames were retained for analysis. MCP foci were detected using the Laplacian of Gaussian (LoG) method. Segmentation and tracking results were visualized in a napari viewer and manually corrected if necessary. MCP states (ON and OFF) were then determined based on the presence of MCP foci in each nucleus, and state transitions were tracked over time. Cumulative counts of nuclei undergoing each transition were calculated.

### Analysis of MCP-GFP and mCherry-PCP movies

Transcriptional foci were identified in images generated by summing the fluorescence intensities from both channels, followed by Gaussian blurring. Spots were detected using the Difference of Gaussians (DoG) method. To quantify the PCP intensity at each detected spot, the mean intensity was measured within a 17×17 pixel square region centered on the spot. Local background intensity was estimated as mean within a larger 31×31 pixel square region surrounding the spot and subtracted from the measured intensity. The detected spots were tracked across frames using the trackpy package, allowing a maximum gap of 5 frames. Tracks that appeared for only a single frame or touched the X-Y borders of the image were discarded. The remaining tracks were sorted based on their time of first appearance in the movie and their total PCP intensity. Montage images were created by cropping a 17×17 pixel square region around each spot in both channels and arranging the images in rows according to tracks and in columns according to time.

### Statistics and reproducibility

All experiments were repeated at least three times with similar outcomes, and representative images are shown in the figures. Two-tailed Mann-Whitney U tests were performed in R (v.3.2.1), followed by Bonferroni correction for multiple comparisons. A value of *P*<0.05 was considered statistically significant. Additional details can be found in the corresponding figure legends.

## RESULTS

### Brd4 bookmarks coincide with early replication

We investigated the role of Brd4 mitotic bookmarks in early *Drosophila* embryos (Fig. 1A). In late S phase 12, Brd4 localizes prominently to histone locus bodies (HLBs), the nuclear bodies associated with the replication-dependent histone gene repeats (Fig. 1B, arrow)(35), as well as to smaller Zelda-dependent foci presumably associated with the zygotically expressed non-histone genes (27). Upon entry into mitosis, Brd4 signals intensify and persist, with dominant HLB-associated clusters localizing near the leading edge of anaphase chromosomes, consistent with their pericentric position and a previous report (36). At the onset of S phase 13, Brd4 clusters rapidly disperse, well before RNA polymerase II clusters appear (Fig. 1B, the 14:40 frame).

**Figure 1.**
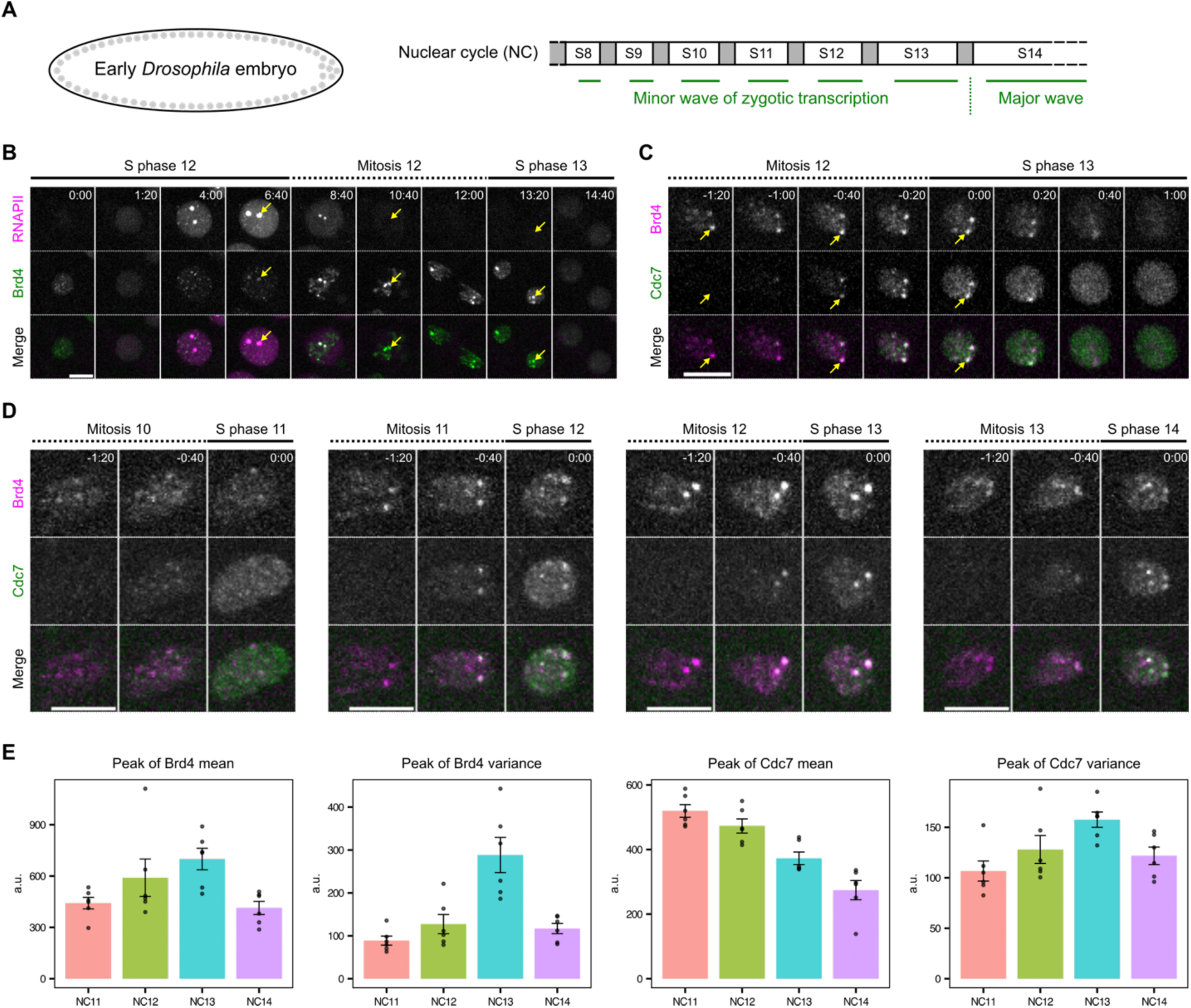
Cdc7-enriched early-replicating hubs emerge concomitantly with the minor wave of zygotic transcription. **A** Schematic of early embryogenesis in *Drosophila*. Following fertilization, the embryo undergoes 13 cycles of synchronous nuclear division cycles, alternating between S phase and M phase (mitosis) without gap phases. A minor wave of zygotic transcription occurs during cycles 8-13, overlapping with the short S phases. DNA replication precedes the onset of transcription after each mitosis, potentially mitigating transcription-replication conflicts during these cycles. The temporal program of DNA replication progressively lengthens from cycle 8 to cycle 13 and is significantly extended in cycle 14, raising the question of how replication and transcription maintain coordination. **B** Representative stills from live imaging of endogenously tagged mCherry-Rpb1, a subunit of RNA polymerase II (RNAPII), and sfGFP-Brd4 in embryos transitioning from early S phase 12 into early S phase 13. The two large RNAPII clusters mark histone locus bodies (arrows), which are assembled at the *HisC* locus encoding ∼100 copies of replication-dependent histone gene array. Brd4 is highly enriched at the *HisC* loci during mitosis 12 and disperses rapidly upon entering S phase 13. **C** Representative stills from live imaging of endogenously tagged mKate2-Brd4 and Cdc7-EGFP in embryos transitioning from mitosis 12 into S phase 13. The S-phase kinase Cdc7 abruptly and transiently forms co-clusters with Brd4 at the *HisC* loci during entry into S phase 13. **D** Representative stills from live imaging of mKate2-Brd4 and Cdc7-EGFP during the transitions from M phase into S phase in successive cycles. All images are maximal projections. Relative time is indicated in minute:second. All scale bars, 5 μm. **E** The peak of either mean or variance of intensities for fluorescently tagged Brd4 or Cdc7 in the nucleus during entry into S phase of indicated nuclear division cycles (NC). Error bars represent SEM, n = 6 embryos.

To test whether Brd4 bookmarks influence replication, we performed two-color live imaging with sfGFP-tagged Brd4 and mCherry-tagged PCNA, a replication fork component that marks replicating loci (12, 37), during the transition from mitosis 12 to S phase 13 (Supplementary Fig. 1A). We observed that mitotic Brd4 clusters began dispersing as the mCherry-PCNA signal appeared in the nucleus at the onset of S phase, with all Brd4 clusters disappearing within 30 seconds. Genomic regions associated with the large HLB-associated Brd4 clusters recruited PCNA faster, resulting in one or two discernible mCherry-PCNA foci in each nucleus about one minute into S phase (Supplementary Fig. 1A, B). To directly test for early replication, we labeled and visualized the earliest replicating DNA in cycles 11-14 by injecting Cy5-dUTP. This revealed foci that were particularly evident in cycles 13 and 14 (Supplementary Fig. 1C), including prominent foci that resembled HLBs as well as other minor foci. These observations document the emergence of early-replicating regions and suggest that these correspond to Brd4-bookmarked regions, particularly the replication-dependent histone genes.

To more directly relate Brd4-bookmarked regions to early replication, we visualized the localization of Cdc7, the catalytic subunit of Dbf4-dependent kinase (DDK) essential for activating pre-replication complexes (pre-RCs). We previously found that endogenously EGFP-tagged Cdc7 begins binding to chromosomes during telophase (38), consistent with its upstream role in initiating replication. Live imaging of mKate2-Brd4 and Cdc7-EGFP showed that Cdc7 rapidly accumulated at HLB-associated Brd4 clusters, alongside a weaker association across chromosomes in late anaphase (Fig. 1C). Within one minute of entering S phase 13, both Brd4 and Cdc7 clusters dispersed and became uniformly distributed in the nucleus. These findings show that a replication initiation factor is transiently enriched at genomic regions bookmarked by Brd4 at the entry into S phase. We refer to the Brd4-Cdc7 co-clusters as “early-replicating hubs”. Notably, we occasionally observed asynchrony in Cdc7 recruitment and dispersal between homologous HLBs within the same nucleus (Fig. 1C), with the slower locus lagging by less than the 20-second frame interval. This asynchrony indicates a lack of temporal coupling between the events at allelic sites.

We conclude that Brd4-bookmarked genomic regions preferentially recruit Cdc7 kinase in late mitosis prior to the onset of S phase. This recruitment facilitates the formation of early-replicating hubs that trigger the onset of replication at select loci.

### Bookmarks and early-replicating hubs change with developmental progression

To investigate developmental regulation, we tracked the dynamics of Brd4 and Cdc7 recruitment during successive nuclear division cycles (Fig. 1D). We focused on the period of initial Cdc7 recruitment, spanning the end of mitosis and the beginning of S phase. From NC11 to NC13, mitotically transmitted Brd4 bookmarks became increasingly prominent, accompanied by intensification of Cdc7 foci at S-phase entry (Fig. 1D). To quantify these changes, we measured the mean and variance of fluorescent intensities for mKate2-Brd4 and Cdc7-EGFP during the 2-minute window following Cdc7 recruitment. We then extracted the peak value from each movie for comparison (Fig. 1E). Brd4 increased in both the peak mean and the peak variance of intensity from NC11 to NC13, consistent with elevated acetylation levels during the minor wave of zygotic transcription. In contrast, Cdc7 decreased in peak mean intensity progressively, suggesting either degradation or titration of Cdc7. However, in parallel with its decrease in peak intensity, the peak variance increased each cycle until NC13, suggesting that, as the maternal supply of Cdc7 becomes limiting with increasing numbers of nuclei, targeted recruitment helps distribute Cdc7 to selected loci and ensure early replication. Interestingly, the trend reversed in NC14, where Brd4 and Cdc7 decreased in both peak mean and variance. This reversal may reflect an increase in early-replicating regions that compete for available Brd4 and Cdc7, ahead of the major wave of zygotic transcription in NC14.

Previous work documented the developmental onset of late replication mediated by the inhibitor Rif1 (12, 38). The observations here suggest a parallel onset of early replication of transcribed regions mediated by Cdc7. Notably, Rif1 and Cdc7 act in opposition to each other in a bistable switch (39), indicating that their balanced activities may shape the global replication-timing program during development.

### CBP-dependent Brd4 bookmarks guide early-replicating hub formation

Given the cell-cycle dynamics of Brd4 clusters (Fig. 1B, C), we hypothesized that events promoting the *de novo* assembly of transcriptional hubs during interphase of one cell cycle (e.g. NC12) generate Brd4 bookmarks that persist through mitosis and are subsequently converted by Cdc7 recruitment into early-replicating hubs in the next cell cycle (e.g. NC13). To test this, we examined whether this cascade depends on CBP, the major lysine acetyltransferase responsible for establishing Brd4 clusters (27).

We first characterized the distribution of endogenously tagged CBP (Fig. 2A, B). During interphase 12, minor foci of mNeonGreen-CBP appeared in a uniform nuclear background, along with enrichment at HLBs evident in late S phase. Upon entry into mitosis 12, CBP dissociated from the chromatin, with the HLB-associated signals disappearing by metaphase. After metaphase, CBP briefly localized to the centrosomes before returning to the nucleus shortly after entry into S phase 13 (Supplementary Fig. 2A). Thus, although CBP is required for the establishment of Brd4 foci (27), it does not maintain an association with these foci during mitosis.

**Figure 2.**
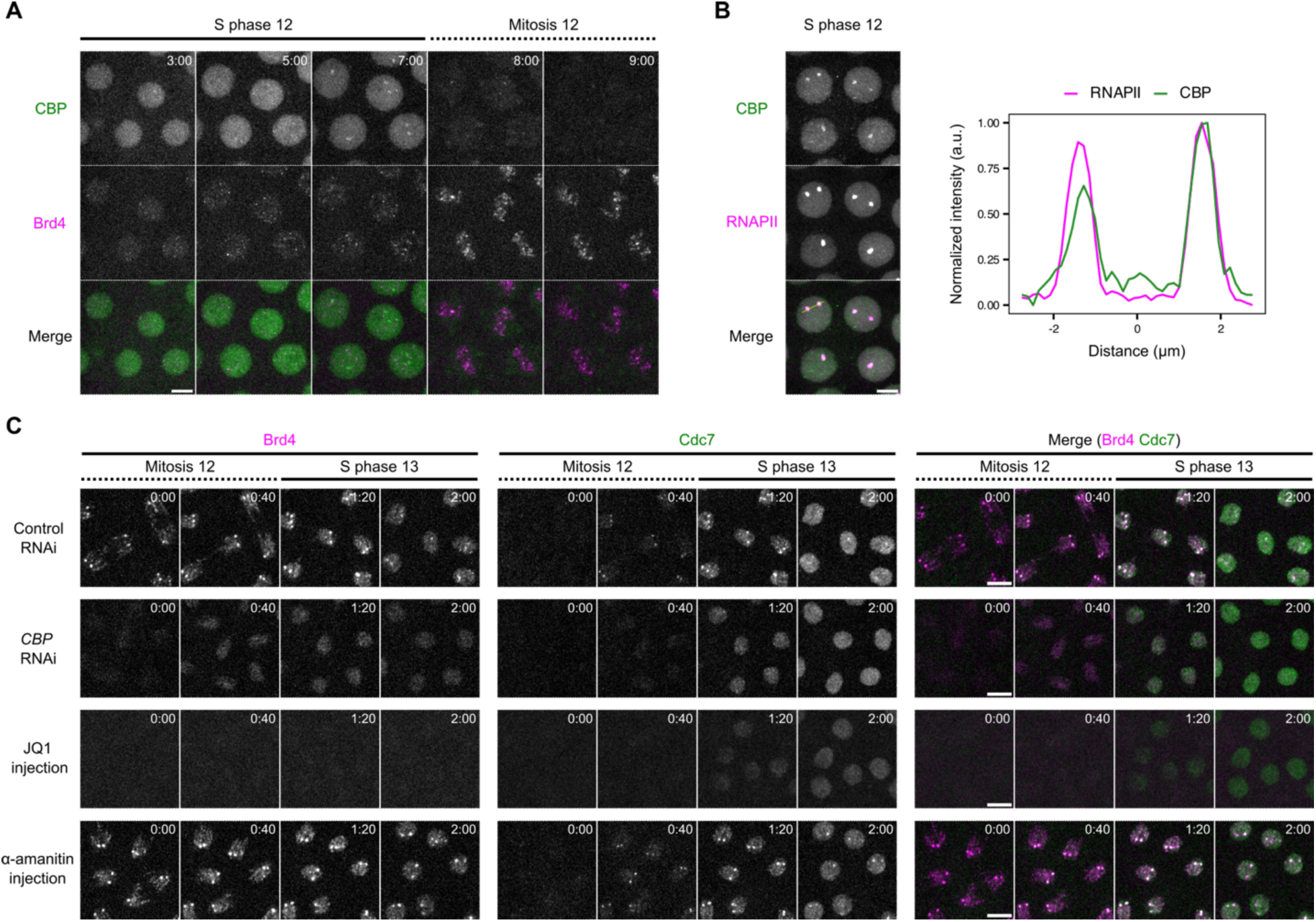
CBP and Brd4 epigenetically link transcriptional activation to the formation of early-replicating hubs across cell cycles. **A** Representative stills from live imaging of mNeonGreen-CBP and mKate2-Brd4 during NC12. Time relative to the start of S phase 12 is indicated in minute:second. Embryos from females heterozygous for both tagged loci (*mKate2-Brd4/+*, *mNeonGreen-CBP/+*) were used for imaging. **B** Snapshot from live imaging of endogenously tagged mNeonGreen-CBP and mCherry-Rpb1 (a subunit of RNAPII) in late S phase 12. Intensity profiles along the line in the merged channel are shown on the right, showing CBP enrichment in the HLBs marked by RNAPII. Embryos from females heterozygous for both tagged loci (*mNeonGreen-CBP/+, mCherry-Rpb1/+*) were used for imaging. **C** Representative stills from live imaging of endogenously tagged mKate2-Brd4 and Cdc7-EGFP in the indicated experiments. The shRNA targeting *white* (*w*) as control or *CBP* was expressed maternally. The Brd4 inhibitor JQ1 and the RNAPII inhibitor α-amanitin were injected around cycle 9-10. All images are maximal projections. Time relative to the start of each movie is indicated in minute:second. All scale bars, 5 μm.

We then examined functional dependency. We used maternally expressed shRNA to knockdown *CBP* in early embryos. As previously reported (27), after *CBP* knockdown, Brd4 clusters at HLBs formed only transiently in interphase. Furthermore, Brd4 did not localize to HLBs on metaphase chromosomes in cycle 12, although weak and uniform binding to anaphase chromosomes still persisted (Fig. 2C and Supplementary Fig. 2C). In addition, CBP depletion delayed nuclear recruitment and abolished the enrichment of Cdc7 at HLBs at the onset of S phase 13 (Fig. 2C and Supplementary Fig. 2C-E). These results support the model that CBP activity establishes Brd4 bookmarks in one cell cycle that persist through the subsequent mitosis to guide the recruitment of Cdc7.

Next, we tested whether Brd4 actively promotes targeted Cdc7 recruitment. We injected the small-molecule inhibitor JQ1 to displace Brd4 from chromatin (Fig. 2C)(40). Loss of Brd4 from mitotic chromosomes delayed the initial recruitment of Cdc7 and abolished its enrichment at HLBs at entry into S phase (Fig. 2C and Supplementary Fig. 2B-E). Thus, Brd4 is required for early and targeted recruitment of Cdc7, rather than simply marking highly acetylated chromatin that replicates early. We conclude that Brd4 is the major effector downstream of CBP to direct the formation of early-replicating hubs at selected loci.

Finally, we asked whether active transcription in earlier cell cycles promotes Brd4 bookmarking. Inhibiting transcription by α-amanitin from cycle 9 to cycle 12 neither reduced Brd4 bookmarking nor blocked Cdc7 enrichment in cycle 13 (Fig. 2C). These findings demonstrate that Brd4 mitotic bookmarks form independently of active transcription and align with previous studies showing that Brd4 recruitment to transcriptional hubs precedes transcription initiation (27).

We conclude that CBP, likely by increasing *de novo* chromatin acetylation at active genes, establishes Brd4 mitotic bookmarks that subsequently promote rapid recruitment of Cdc7 kinase prior to the onset of S phase, providing one mechanism that links transcriptional activation in one cycle to early replication in the next.

### Cdc7 activity and the ephemeral existence of early-replication hubs

Given that the replication licensing mechanism limits replication to once and only once per cell cycle, early-replicating hubs would not be able to trigger a second round of replication in the same cycle. However, if localized Brd4 persistently promoted transcription, it would clash with the locally induced replication. Consequently, the subsequent fate of early-replicating hubs is important. As shown above, Brd4-Cdc7 co-clusters disperse within a minute after emergence (Fig. 1C). We hypothesized that localized Cdc7 activity, or a downstream event such as replication initiation, triggers their disassembly. To test this, we injected embryos with the Cdc7 inhibitor XL413 (Cdc7i) and used live imaging to examine the effect on early-replicating hubs during entry into S phase 13 (Fig. 3A, B and Supplementary Fig. 3A).

**Figure 3.**
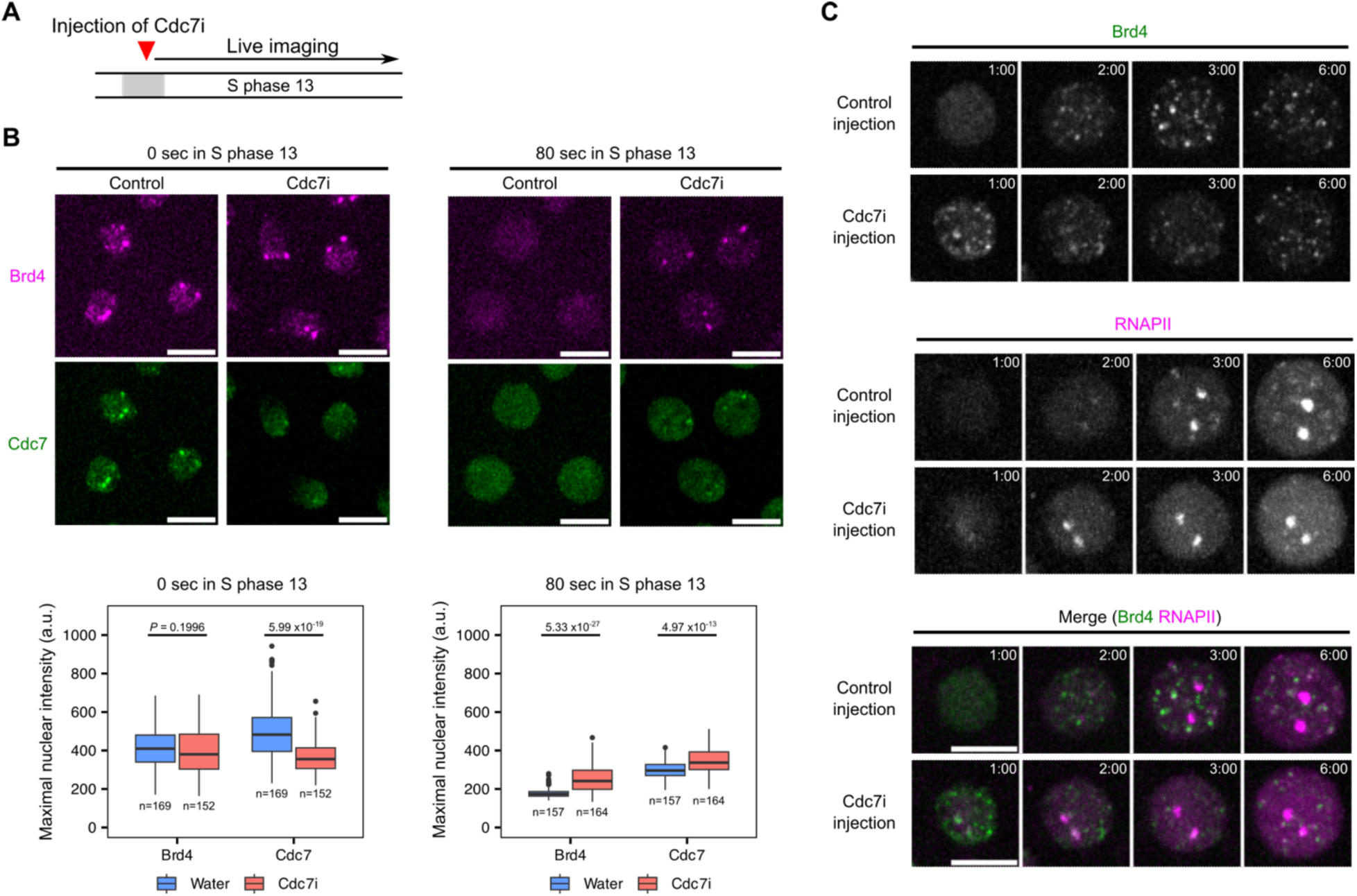
Cdc7 controls its transient localization at early-replicating hubs, erases Brd4 bookmarks, and delays RNAPII recruitment. **A** Experimental design for determining whether Cdc7 activity influences the formation or dispersal of early-replicating hubs. Either water (control) or XL413, a Cdc7 inhibitor (Cdc7i), was injected into embryos, followed by live imaging during S phase 13. The injection of Cdc7i was carried out during mitosis 12 to avoid cell-cycle defects prior to S phase 13. **B** (top), Snapshots from live imaging of mKate2-Brd4 and Cdc7-EGFP in control or Cdc7i-injected embryos at the indicated times in S phase 13. **B** (bottom), Box plots showing the maximal intensities for fluorescently tagged Brd4 or Cdc7 in nuclei at the indicated times. The central lines of the box plots represent median. Whiskers extend to 1.5 times interquartile range from the box. The number of nuclei (*n*) pooled from 3 embryos were indicated. *P* values were calculated using two-sided Mann-Whitney U test with Bonferroni correction for multiple comparisons. a.u., arbitrary unit. **C** Representative stills from live imaging of sfGFP-Brd4 and mCherry-Rpb1 (a subunit of RNAPII) in control or Cdc7i-injected embryos. All images are maximal projections. Time relative to the start of S phase 13 is indicated in minute:second. All scale bars, 5 μm.

In control embryos injected with water, Cdc7-EGFP formed prominent co-clusters with the mKate2-Brd4 at HLBs at the beginning of S phase (Fig. 3B, left; Supplementary Fig. 3A). In contrast, Cdc7i injection disrupted targeted Cdc7 recruitment, resulting in uniform localization of Cdc7 in the nucleus, despite the presence of Brd4 bookmarks (Fig. 3B, left; Supplementary Fig. 3A). This shows that Cdc7 activity contributes to its targeted recruitment to Brd4-bookmarked regions, suggesting a positive feedback loop potentially generating a switch-like behavior. Stochastic transition of such a switch could explain the slight asynchrony in Cdc7 recruitment between homologous HLBs.

Additionally, Brd4 bookmarks persisted longer in Cdc7i-injected embryos compared to the rapid dispersal in control embryos (Fig. 3B, right; Supplementary Fig. 3A). While reduced, a low level of localized Cdc7 persisted at Brd4 foci in these inhibitor injected embryos. This low level of localized Cdc7 persisted beyond than that in control embryos (Fig. 3B, right; Supplementary Fig. 3A). Using maximal fluorescent intensity as a proxy for protein clustering, we confirmed a decrease in initial Cdc7 enrichment, along with prolonged presence of Brd4 and Cdc7 clusters in Cdc7i-injected embryos (Fig. 3C). These findings suggest that Cdc7 activity plays a dual role in early-replicating hubs: initially facilitating its own recruitment to Brd4-bookmarked regions to form hubs and subsequently promoting hub disassembly marked by rapid dispersal of both Brd4 and Cdc7 foci.

Given the known role of Brd4 in recruiting RNAPII and promoting transcription (27), we wondered whether persisting Brd4 bookmarks in Cdc7i-injected embryos were associated with dysregulated transcription. To test this, we injected Cdc7i into embryos expressing sfGFP-tagged Brd4 and mCherry-tagged Rpb1 (the largest subunit of RNAPII) and performed live imaging during S phase 13. In control embryos, Brd4 bookmarks dispersed completely by 1 minute after S phase entry but began to re-emerge at 2 min, followed by RNAPII recruitment to HLBs (Fig. 3C). In Cdc7i-injected embryos with persisting Brd4 bookmarks, RNAPII foci emerged earlier within 2 minutes into S phase. Repeating this experiment in embryos expressing mCherry-PCNA confirmed that Cdc7i blocked the formation of early PCNA foci while advancing the recruitment of RNAPII to HLBs (Supplementary Fig. 3B). Despite advanced recruitment, RNAPII levels at HLBs did not reach those observed in control embryos later in S phase (Supplementary Fig. 3C), possibly due to underreplicated DNA templates or conflicts between transcription and replication machinery.

We conclude that Cdc7 activity promotes timely removal of Brd4 bookmarks and the disassembly of early-replicating hubs. This chromatin resetting delays RNAPII recruitment, ensuring the coordination such that onset of transcription follows onset of replication.

### Cdc7 coordinates transcriptional states with S-phase entry

We previously reported that DNA replication is essential for transcriptional hub formation and the onset of transcription in NC12 (11). This contrasts with the above data suggesting that an inhibitor of DNA replication, Cdc7i, advances RNAPII recruitment in NC13. Given the finding that inhibition of Cdc7 stabilizes pre-existing Brd4 foci, we hypothesized that abnormal retention of Brd4 bypassed the need for *de novo* transcriptional hub formation, and that this bypass predominated in cycles 13 and 14. To explore this, we re-examined the effects of replication inhibitors on RNAPII dynamics during successive nuclear cycles (Fig. 4A).

**Figure 4.**
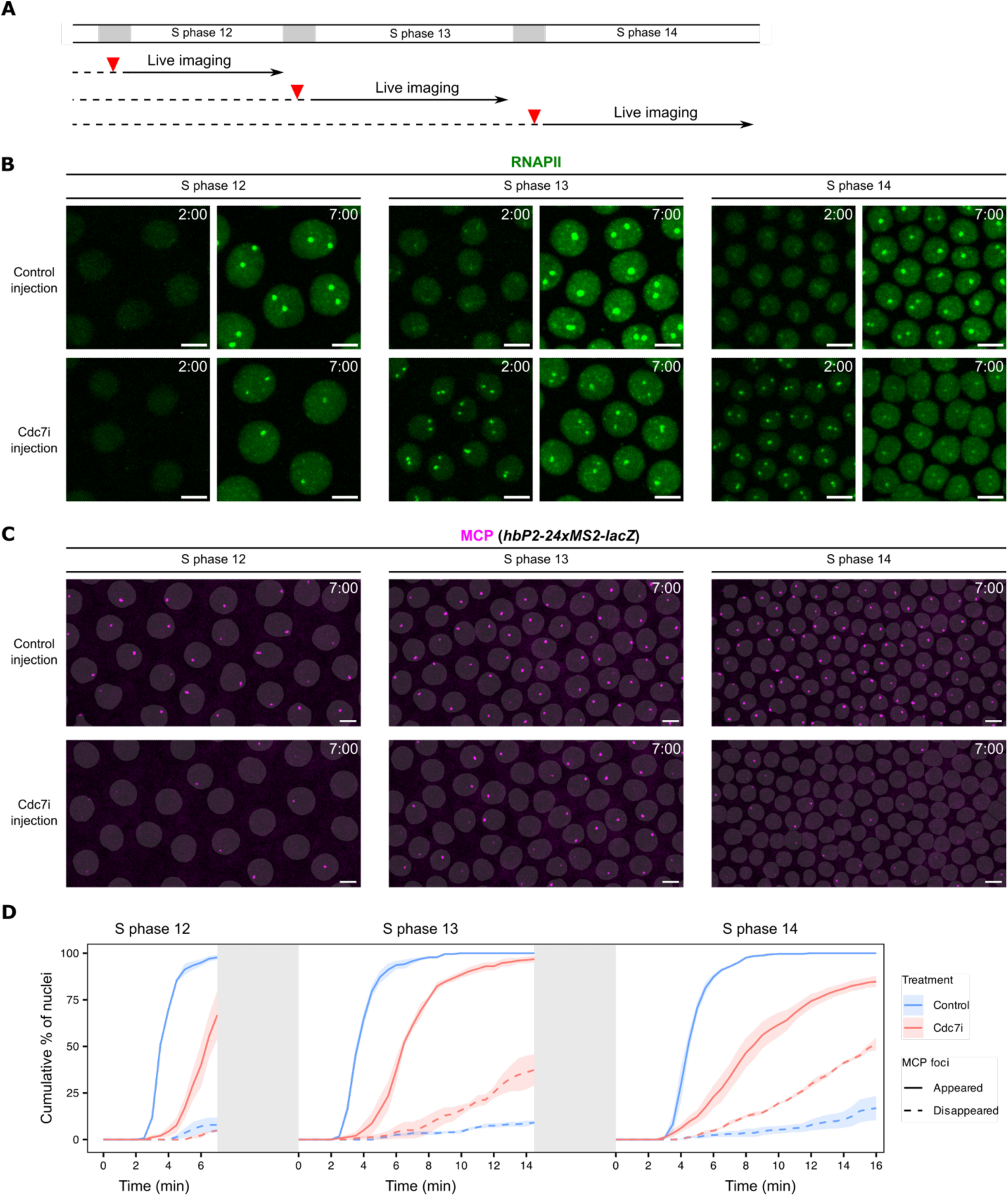
Cdc7 activity delays and synchronizes RNAPII recruitment and prevents precocious termination of transcription during cycles 13 and 14. **A** Experimental design for determining the impact of Cdc7 inhibition on transcription in different nuclear cycles. Cdc7 inhibitor (Cdc7i) was injected into staged embryos during mitosis, followed by live imaging in the subsequent S phase. Control embryos were injected with water during the preceding S phase or mitosis. **B** Representative snapshots from live imaging of EGFP-Rpb3 (a subunit of RNAPII) in control or Cdc7i-injected embryos at the indicated times during S phase of various cycles. **C** Representative snapshots from live imaging of MCP-mCherry in control or Cdc7i-injected embryos with a *hbP2-24xMS2-lacZ* reporter. Frames at 7 minutes in S phase are shown. The binary masks of nuclei were generated from EGFP-Rpb3 images and overlaid on the MCP-mCherry images. Time relative to the start of S phase is indicated. All images are maximal projections. All scale bars, 5 μm. **D** Cumulative percentage of nuclei that have acquired MCP foci (“MCP turned ON”) and the cumulative percentage that subsequently lost MCP foci (“MCP turned OFF”) since entering an S phase following control or Cdc7i injection. Colored shading represents SEM, n = 3 embryos. The grey blocks represent the interleaving mitosis and do not report actual duration.

First, to see whether the change in response to Cdc7 inhibition reflected a general change in the coupling of transcription to replication, we injected Geminin into NC13 embryos to block origin licensing with a resulting block to replication in NC14. This treatment strongly inhibited RNAPII recruitment to HLBs in NC13 (Supplementary Fig. 4A), consistent with the effects observed in NC12 and NC13 (11). Based on the known action of Geminin in blocking pre-replication complex formation and thus replication, we suggest that this is consistent with a coupling of transcription to replication, although it is possible that Geminin acts via a different route to inhibit RNA polymerase recruitment.

Next, we confirmed our previous finding that Cdc7i injection strongly delayed the recruitment of RNAPII to histone genes during NC12, resulting in significantly smaller HLB-associated RNAPII clusters that only formed in late S phase (Fig. 4B)(11). However, in both NC13 and NC14, Cdc7i injection advanced RNAPII recruitment, leading to more prominent HLBs at 2 minutes into S phase compared to control embryos (Fig. 4B). These prematurely recruited RNAPII clusters were unstable and diminished rapidly within 5 minutes, whereas those RNAPII clusters in control embryos continued to grow substantially (Fig. 4B). These results suggest that Cdc7 has multiple roles in coordinating transcription and replication: fulfilling a requirement for transcriptional hub assembly in earlier cycles, while preventing the premature RNAPII recruitment to bookmarked genes in later cycles.

Given the premature but unstable recruitment of RNAPII to histone genes upon inhibition of Cdc7, we next examined how Cdc7 inhibition impacts the onset of transcription. For this live-imaging analysis, we used a *hunchback* (*hb*) transcriptional reporter gene (10), which is broadly expressed in the anterior half of the embryo from NC9 onwards. We used embryos maternally expressing MCP-mCherry and carrying a *hbP2-MS2-lacZ* construct. In control embryos, MCP foci appeared in all nuclei in the anterior domain, and the majority of these foci persisted stably (Fig. 4C). Injection of Cdc7i led to irregularities in both the activation of expression and an instability in the maintenance of expression. These irregularities became more predominant as S-phase duration increased in successive NCs (Fig. 4C). In some nuclei, faint MCP foci appeared and then disappeared within 2 minutes (Supplementary Fig. 4B). To quantify these effects, we measured the cumulative percentage of nuclei in which MCP foci appeared, as well as those in which MCP foci appeared and subsequently disappeared after entering S phase (Fig. 4D). In control embryos, MCP foci appeared synchronously within 5 minutes after each mitosis, and the majority of them persisted throughout the imaging window. In Cdc7i-injected embryos, MCP foci appeared more slowly, and an increased number of them disappeared early. Because the MS2 cassette is inserted in the 5’UTR of this reporter, this short lifetime suggests either a reduced transcriptional burst size or premature transcript abortion upon Cdc7i injection.

We conclude that Cdc7 kinase activity delays RNAPII recruitment in later cycles, contributes to the replication-first ordering, and stabilizes transcriptional state potentially through minimizing collisions.

### Early Cdc7 activity safeguards transcriptional elongation

To further investigate the cause of premature transcriptional termination in Cdc7i-injected embryos, we used a modified *hunchback* transcriptional reporter construct containing MS2 repeats in the 5’UTR and PP7 repeats in the 3’UTR (41). This dual labeling allowed us to monitor both transcriptional initiation and elongation in real time (Fig. 5A).

**Figure 5.**
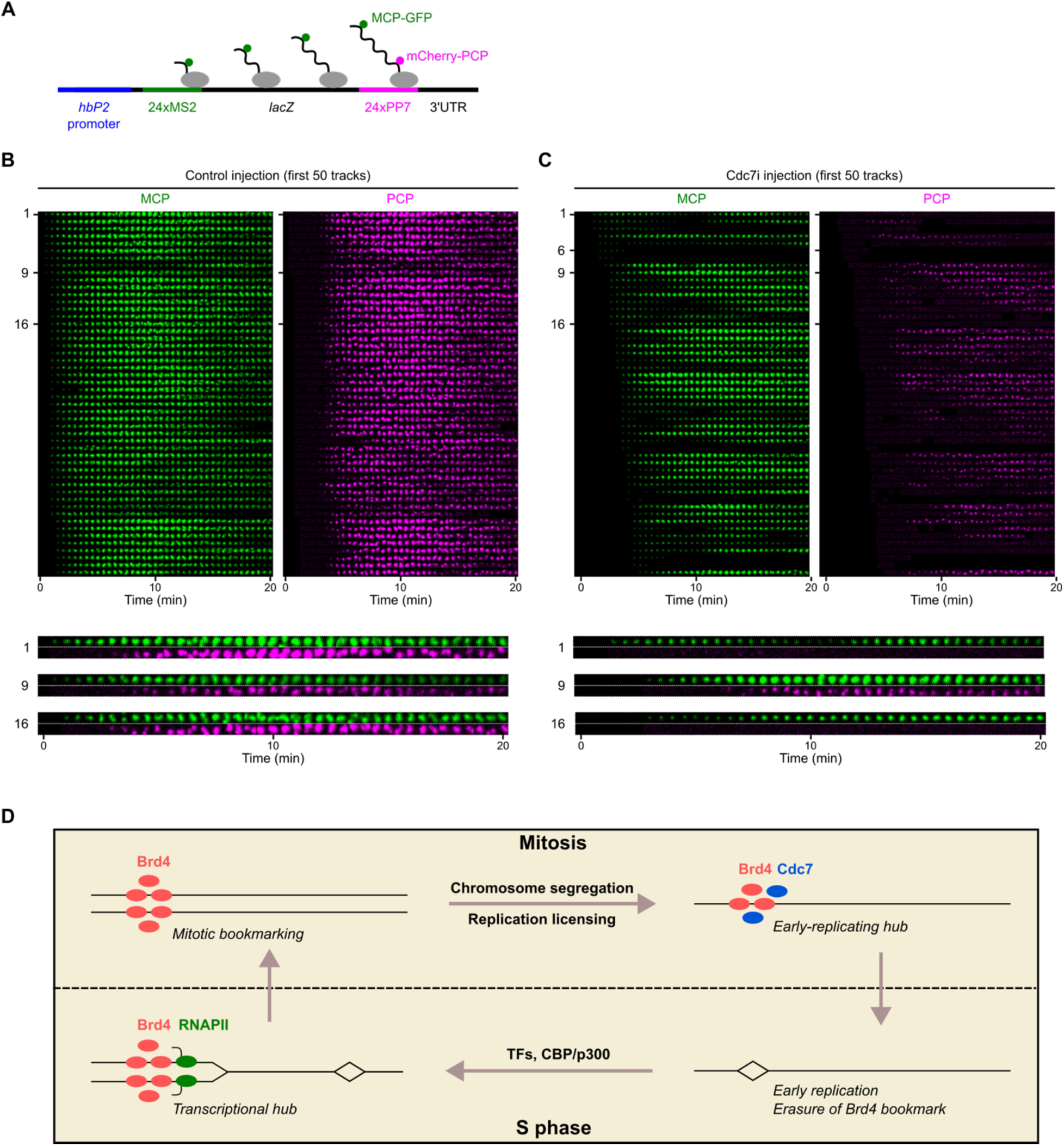
Early replication safeguards transcriptional elongation at the midblastula transition. **A** Schematic of the *hbP2* transcriptional reporter that contains 24x MS2 loops in the 5’UTR and 24x PP7 loops in the 3’UTR, enabling visualization of the transcripts by MCP-GFP and mCherry-PCP, respectively. **B, C** Montages with images in each row representing the signal from a single reporter locus as it changes over time in control **(B)** or Cdc7i-injected embryos **(C)** during S phase 14. The top 50 spots with the earliest MCP appearance from a representative embryo in each group are displayed. Similar outcomes were observed in three embryos. The rows are ordered first by MCP emergence time and then by PCP intensity. Time relative to the initial appearance of MCP foci in the same embryo is indicated. Magnified images of selected spots are shown at the bottom. **D** Model for the coordination of replication and transcription by Brd4 bookmarks in early *Drosophila* embryo.

In control NC14 embryos, MCP foci marking the 5’ MS2 appeared synchronously, consistent with rapid onset of transcription (Fig. 5B). These MCP foci were soon joined by PCP foci marking the 3’ PP7 sequence after 3-4 minutes, indicating normal transcript elongation. In contrast, Cdc7i-injected NC14 embryos displayed a range of transcriptional defects, including delayed onset of transcription, reduced transcript levels, and impaired elongation (Fig. 5C and Supplementary Fig. 5). Specifically, we observed the following defects with regard to elongation: (1) transient MCP foci that disappeared without acquiring PCP foci (track 6); (2) unstable MCP foci that fluctuated in intensity but did not acquire PCP foci (track 1); (3) stable MCP foci that acquired PCP with a delay and at lower levels (track 9); and (4) stable MCP foci that never acquired PCP throughout live imaging (track 16). Notably, we did not observe a clear relationship between the severity of defects and the distance from the injection site (Supplementary Fig. 5A), suggesting that the variability in response was not due to local differences in Cdc7i concentration. Since Cdc7i only reduces the probability of pre-RC activation without completely inhibiting it, the stochastic nature of active replication fork production could result in the observed heterogeneity in transcription defects.

We conclude that Cdc7 activity, likely by advancing local onset of replication, coordinates replication and transcription to avoid collisions that disrupt transcriptional elongation as the duration of S phase increases in the last two syncytial cycles (NC13 and NC14).

## DISCUSSION

How cells coordinate transcription and replication to prevent conflicts remains an open question, with broad implications for genome integrity. In previous work, we proposed a collision-avoidance mechanism in early *Drosophila* embryos, wherein transcription occurs only on newly replicated DNA during each S phase, allowing transcripts to elongate behind replication forks unimpeded (11). This temporal ordering was shown to be enforced in cycle 12 by making the onset of transcription dependent on replication. Accompanying developmental changes in the cell cycle, particularly the emergence of a replication-timing program and the lengthening of S phase, a second mechanism comes into play in cycle 13 to reinforce the temporal ordering (Fig. 5D). This mechanism is based on the control of replication timing by a mitotic bookmark at transcriptionally activated genes. By advancing the replication of transcriptionally active regions, the bookmark prepares the region for subsequent transcription without the risk of collisions.

This bookmarking mechanism involves the transcriptional coactivator Brd4. We show that Brd4, in addition to localizing to transcriptional hubs and promoting transcription in S phase 12 (27), has a second function in bookmarking specific genes on mitotic chromosomes (Fig. 5D). Towards the end of mitosis and the onset of S phase 13, Brd4 bookmarks directly or indirectly recruit the replication initiation factor Cdc7. Thus, Cdc7 repurposes the bookmark to stimulate early replication instead of transcription (Fig. 5D). In addition to promoting local replication, the dispersal of Brd4 bookmarks by Cdc7 eliminates its potential to promote transcription, thereby enforcing a requirement for *de novo* assembly of a new transcriptional hub to start a new cycle of transcription and gene bookmarking. Notably, this dependency on the formation of a new transcriptional hub allows transcription factor reprograming and delays transcription so that it follows onset of replication of the locus. These findings reveal that, as embryogenesis progresses and cell-cycle regulation matures, embryos introduce additional measures to temporally order replication and transcription, thereby avoiding collisions.

Mitotic bookmarking of selected loci by transcription factors or chromatin regulators has been associated with rapid transcriptional reactivation in a subsequent cell cycle (26). In this study, we uncover an additional role for mitotic bookmarks in promoting early replication of bookmarked loci in embryonic cycles that lack a G1. Notably, *Drosophila* embryos exhibit transcriptional memory (42, 43), in which reporter genes stochastically expressed in one cycle tend to be reactivated more rapidly in subsequent cycles. Our findings herein suggest that this transcriptional memory may arise indirectly from advanced replication timing. That is, by accelerating replication of transcriptionally active regions, bookmarks indirectly establish an early competence for collision-free transcription upon mitotic exit.

We emphasize that our findings show that the action of Cdc7 is responsible for repurposing the use of bookmark, thus the timing of this repurposing ought to depend on cell-cycle regulation of Cdc7 activity. Cdc7 kinase with its regulatory subunit, Dbf4 in budding yeast and Chiffon in *Drosophila*, forms an active kinase complex called DDK (Dbf4-dependent kinase). DDK together with the cyclin-dependent kinase (CDK) phosphorylate the pre-replication complex (pre-RC) during S phase and initiate the assembly of bi-directional replication forks. In a mature cell cycle, DDK and S-phase CDK are only activated at the G1-S transition, suggesting that the repurposing of bookmark function from transcription to replication is deferred until the onset of S phase in cells with a G1 phase. Indeed, we show that the reduction in Cdc7 activity in early *Drosophila* embryo is sufficient to allow persistence of the Bdr4 bookmark that retains an ability to recruit RNAPII and activate transcription. These results suggest that Brd4 bookmarks have the potential of promoting post-mitotic transcription in a cell cycle with G1, while in the embryonic cell cycles, highly abundant DDK and CDK override this potential by triggering early replication. Hence, though our characterization is currently limited to the early embryonic cycles of *Drosophila*, we propose that Cdc7 acts widely to repurpose the bookmark activity from transcription to replication, and that the dual use of the bookmark underlies the widely observed finding that transcriptionally active regions replicate early (14).

Since the Brd4 mitotic bookmark is bifunctional, it seemingly has the potential to activate both replication and transcription at the same time and induce collisions. To avoid this, the bookmark has to maintain exclusivity, such that triggering replication is accompanied by inhibiting transcription, and vice versa, so that it never does both at the same time. In part, this exclusivity appears to be due to a dual action of Cdc7: the recruitment of Cdc7 to the Brd4 bookmark both activates pre-RCs to promote early replication and disperses Brd4 bookmarks to prevent immediate transcription. But, with a time lag of about 2 min, new transcriptional hubs are built: what prevents the newly produced hubs from activating replication as well as transcription. We suggest that the replication licensing system prevents the newly acquired Brd4 bookmarks from reactivating replication. DNA replication is regulated so that all sequences replicate once and only once in a typical cell cycle. The license to replicate requires the assembly of the pre-RC during cell-cycle stages in which kinases activating the pre-RC are low (anaphase in the case of early nuclear cycles of syncytial *Drosophila* embryo). Following licensing, replication only occurs after the activation of pre-RCs by S-phase kinases. Once activated, pre-RCs cannot reform until the passage of another cell cycle. When a new Brd4 bookmark is deposited on already replicated DNA, it cannot promote replication without the formation of a new pre-RC, and thus it is limited to promoting transcription until re-licensing occurs during the next mitosis in nuclear division cycles or in late G1 in more mature cell cycles. In this way, the DNA licensing system further enforces the exclusivity of Brd4 bookmark function, which we propose coordinates transcription and replication timing to avoid collisions.

Our study also provides insights into how the replication-timing program emerges and what function it serves during embryonic development. Previous work showed that transcription factors, acting through lysine acetyltransferases, promote the deposition of a Brd4 bookmark that acts during one cell cycle to promote locus-specific transcription (27). We now show that this mark persists into the next cell cycle where it locally promotes the early replication of licensed DNA sequences. The formation of these hubs enriched in Cdc7 kinase depends on the action of the lysine acetyltransferase CBP in the previous cell cycle. Given the conserved roles of CBP/p300 in transcriptional activation across species (28, 29, 44, 45), we suggest that these mechanisms are used widely to link the switching of transcriptional state in one cell cycle to the switching of replication timing in the next cycle. Our data demonstrate that early replication has a protective role for transcription. This mechanism may be especially important for genes transcribed in early S phase, such as replication-dependent histone genes. We suggest that a replication-first strategy broadly serves to coordinate transcription and replication to protect genome integrity.

Previous work identified several strategies that coordinate transcription and replication. Microscopy revealed spatial separation of transcription and replication sites throughout the nucleus even within nucleoli (8, 46, 47). Such a separation requires controls coordinating the two processes. Since all DNA sequences must replicate, avoidance of overlap requires a pause in transcription during the replication of transcribed sequences. Numerous mechanisms have been suggested to contribute to this type of coordination. Transcription can be transiently shut down ahead of replication forks by a yet unclear mechanism (6, 7), and stalled RNAPII may be removed by fork-associated Integrator complexes (48). Conversely, transcription can influence the initiation sites and progression of replication to minimize conflicts within actively transcribed genes (49, 50). Finally, replication fork barriers, such as those in rDNA loci, help prevent replication forks from invading transcribed regions (51). Our findings expand this repertoire of strategies by showing how a mitotic bookmark guides early replication to safeguard S-phase transcription. This opens up new opportunities to dissect the precise crosstalk between transcription and replication, which is critical for preserving transcriptional fidelity and genome stability.

## ACKNOWLEDGEMENTS

Stocks obtained from the Bloomington Drosophila Stock Center (NIH P40OD018537) were used in this study. We thank Takashi Fukaya for sharing the fly lines to visualize nascent transcripts containing 5’ MS2 and 3’ PP7 repeats. We thank Ekaterina Korotkevich and Robert J. Duronio for helpful comments on the manuscript and Barbara Panning for discussion.

## AUTHOR CONTRIBUTIONS

**C.-Y.C.** Conceptualization, Methodology, Formal analysis, Investigation, Writing – Original Draft, Visualization.

**P.O’F**: Conceptualization, Writing – Original Draft, Supervision, Funding acquisition.

## DATA AVAILABILITY

Raw imaging data generated in this study have been deposited at Zenodo (https://zenodo.org/records/13761202). Custom code used for data analysis and visualization is available on GitHub (https://github.com/ccho3/bookmarking-DNA-replication). Any additional information is available from the corresponding author upon request.

