## Supplementary Information for "A mitotic bookmark coordinates transcription and replication"

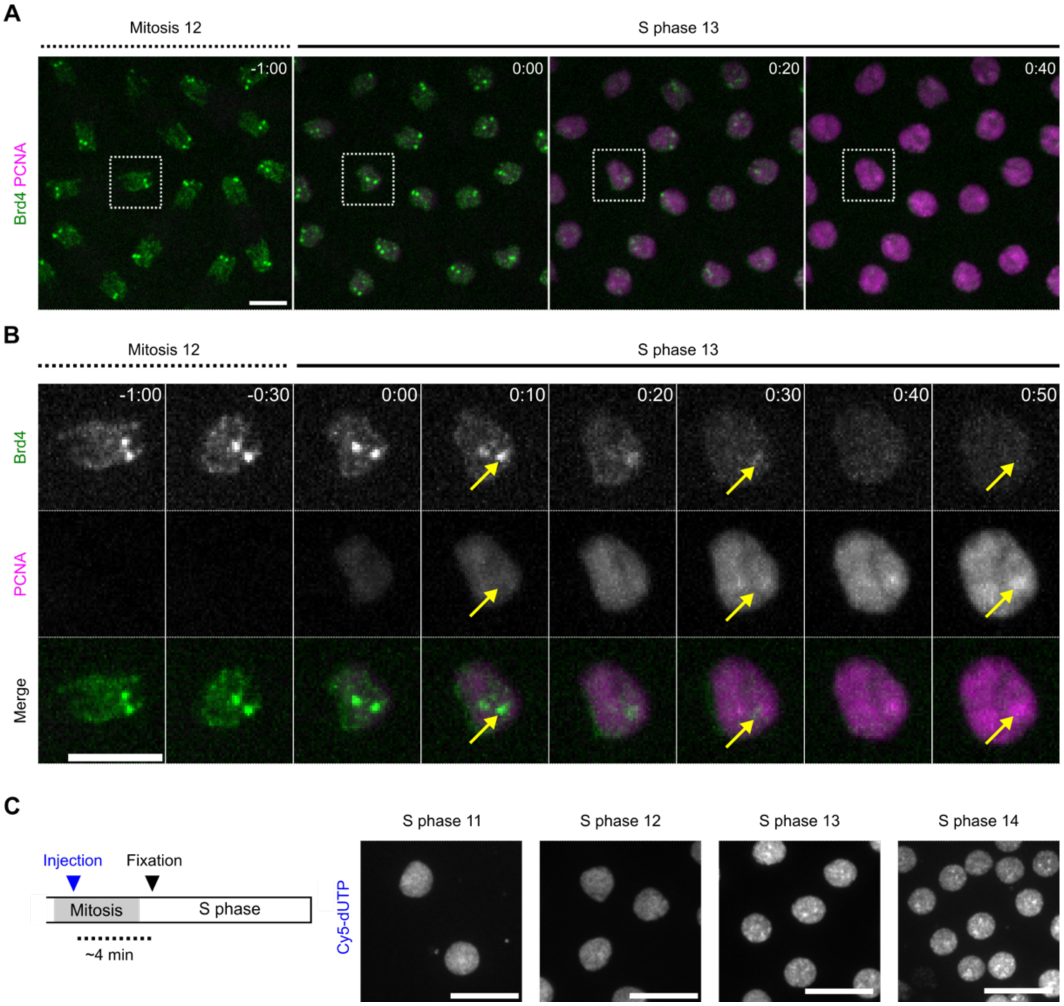


**Supplementary Figure 1. Additional data related to Figure 1. A, B** Representative stills from live imaging of endogenously tagged sfGFP-Brd4 and transgenic mCherry-PCNA in embryos transitioning from mitosis 12 into S phase 13. The boxes in panel **A** outline the same nucleus shown in panel **B**. **C** (left), Experimental design for labeling early-replicating DNA. A pool of unsynchronized embryos were injected with Cy5-dUTP and fixed after 4 minutes, and then those embryos that just entered S phase were identified by the relatively compact chromatin and small nuclear size. Given that mitosis in the early embryo takes approximately 5 minutes, labeling of late-replicating sequences from the previous cycle was effectively prevented. **C** (right), Representative images of fixed embryos subjected to short pulse-labeling with Cy5-dUTP at the beginning of S phases. Prominent early-replicating domains emerge after S phase 13.

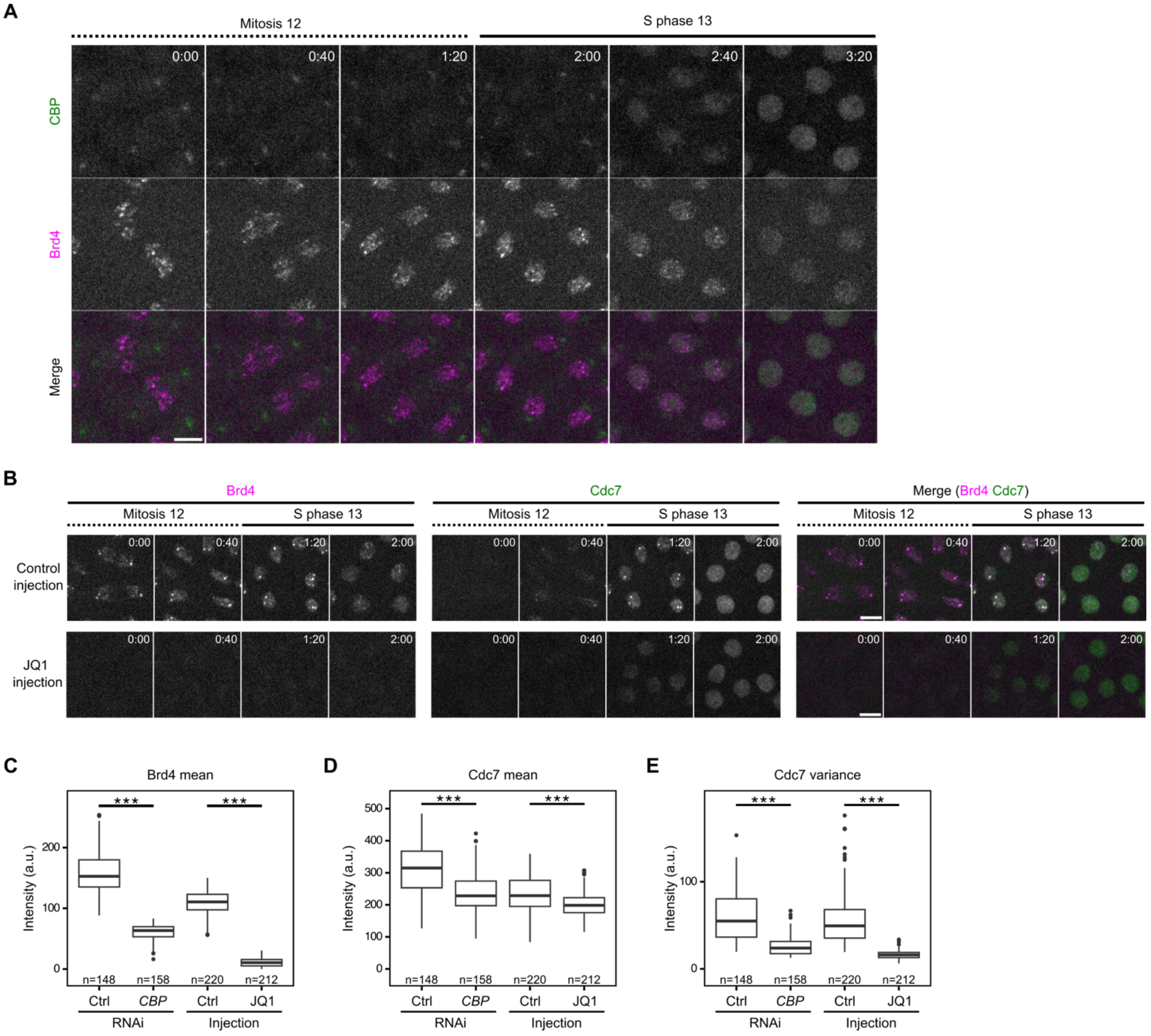


**Supplementary Figure 2. Additional data related to Figure 2. A** Representative stills from live imaging of mNeonGreen-CBP and mKate2-Brd4 during the transition from mitosis 12 into S phase 13. **B** Representative stills from live imaging of endogenously tagged mKate2-Brd4 and Cdc7-EGFP in embryos injected with 1% DMSO (control) or the Brd4 inhibitor JQ1. All images are maximal projections. Time relative to the start of each movie is indicated in minute:second. All scale bars, 5 μm. **C-E** Box plots showing the mean or variance of intensities for fluorescently tagged Brd4 or Cdc7 in nuclei at the start of S phase 13 in the indicated experiments. The central lines of the box plots represent median. Whiskers extend to 1.5 times interquartile range from the box. The number of nuclei (*n*) pooled from 5 embryos were indicated. ***, $P < 0.001$ by a two-sided Mann-Whitney U test with Bonferroni correction for multiple comparisons. a.u., arbitrary unit.

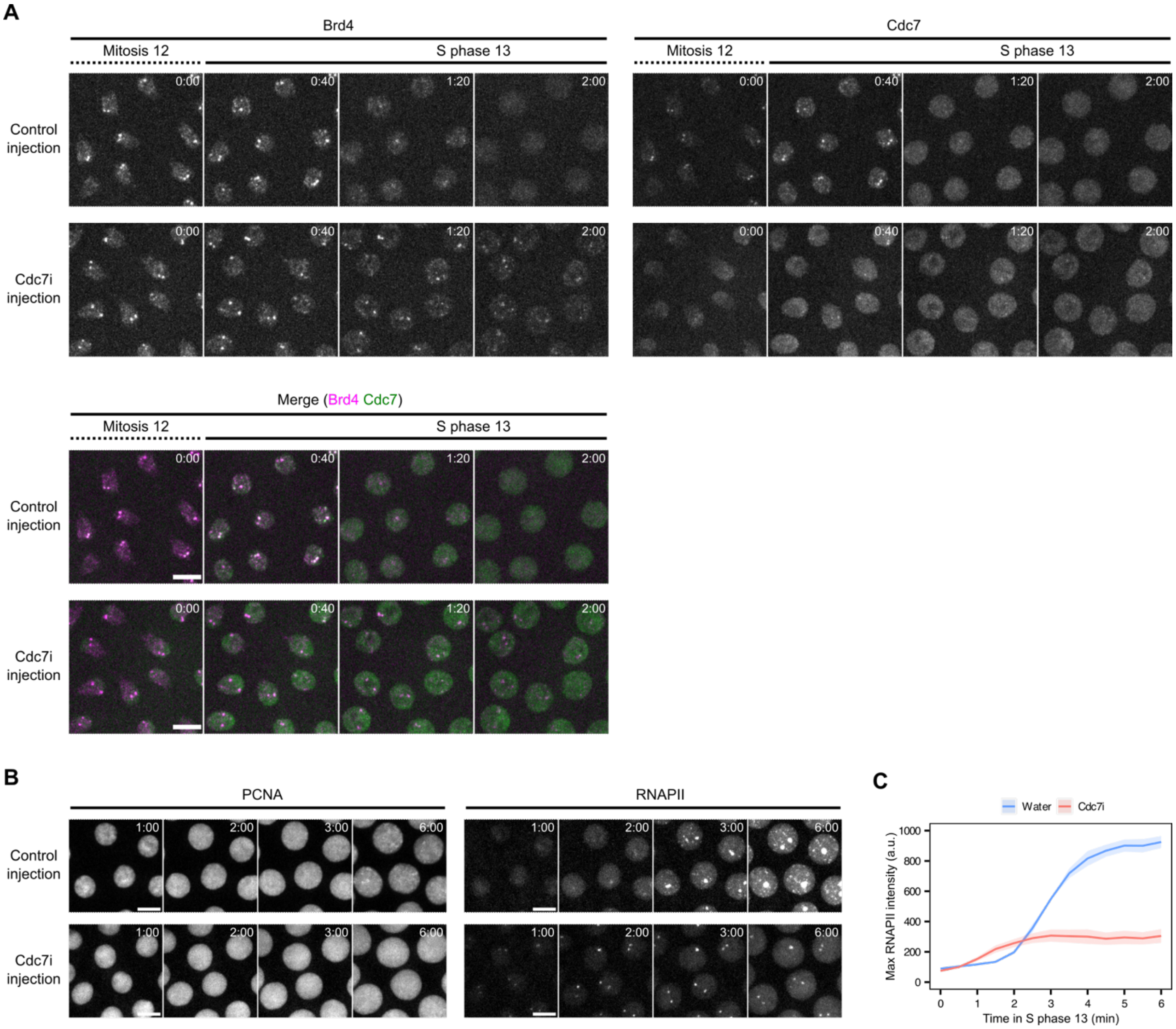


**Supplementary Figure 3. Additional data related to Figure 3. A** Representative stills from live imaging of endogenously tagged mKate2-Brd4 and Cdc7-EGFP in embryos injected with water (control) or Cdc7 inhibitor (Cdc7i) and transitioning from mitosis 12 into S phase 13. Time relative to the start of the movie is indicated in minute:second. **B** Representative stills from live imaging of mCherry-PCNA and EGFP-Rpb3 (a subunit of RNAPII) in control or Cdc7i-injected embryos during S phase 13. Recombinant mCherry-PCNA was co-injected with control or Cdc7i. Time relative to the start of S phase is indicated in minute:second. All images are maximal projections. All scale bars, 5 $\mu$m. **C** Quantification of nuclear EGFP-Rpb3 maximal intensity in control or Cdc7i-injected embryos during early S phase 13. The maximal intensity was measured in all nuclei and averaged per embryo. Shaded areas represent SEM, n = 3 embryos.

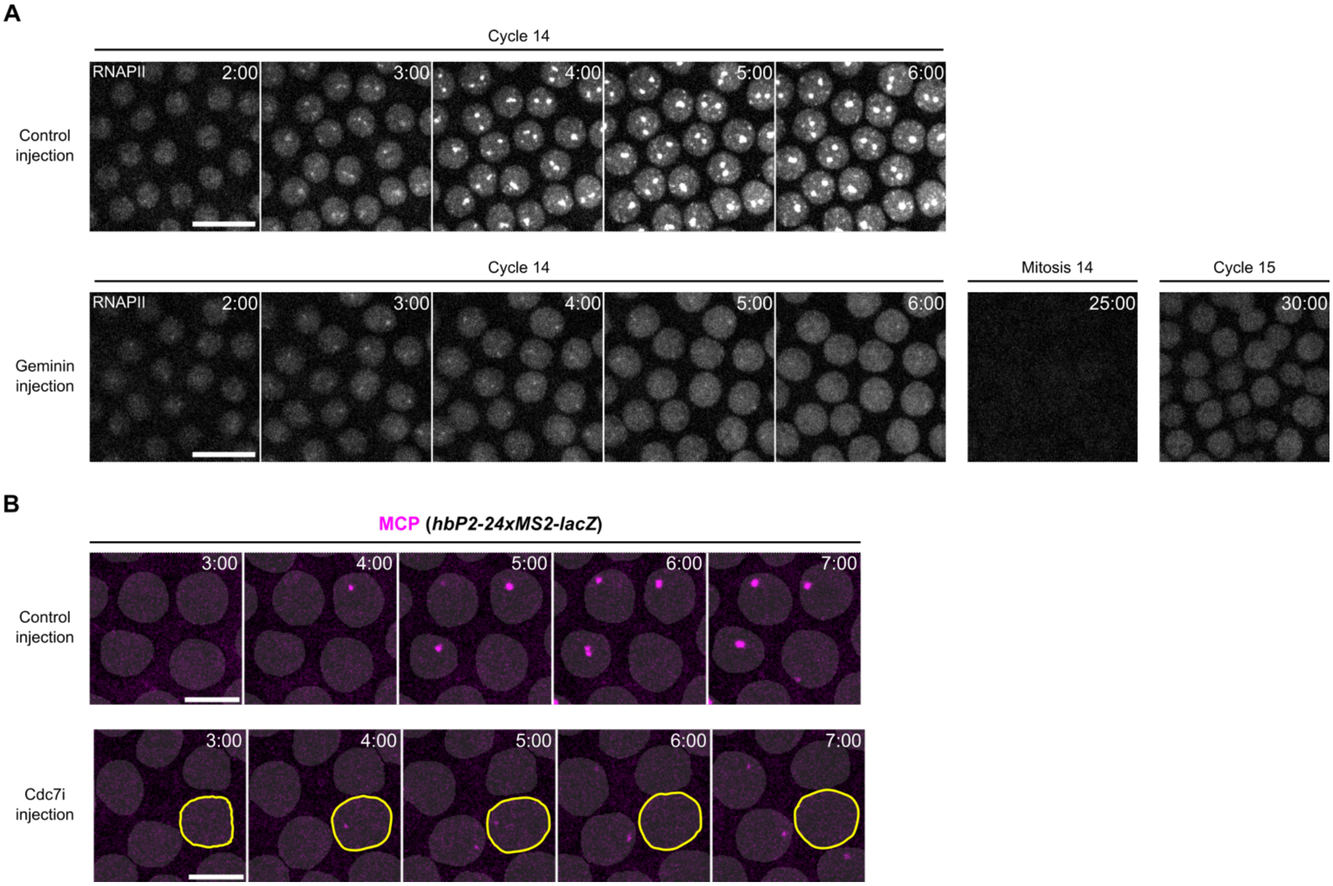


**Supplementary Figure 4. Additional data related to Figure 4. A** Representative stills from live imaging of mCherry-Rpb1 (RNAPII) in control or Geminin-injected embryos in S phase 14. The injection was performed in S phase 13. Time relative to the start of S phase 14 is indicated in minute:second. The Geminin-injected embryo underwent an extra cycle of synchronous division as a result of S-phase ablation. Scale bars, 10 μm. **B** Representative stills from live imaging of MCP-mCherry in control or Cdc7i-injected embryos with a *hbP2-24xMS2-lacZ* reporter. The binary masks of nuclei were generated from EGFP-Rpb3 images and overlaid on the MCP-mCherry images. Yellow circles highlight a nucleus whose MCP foci emerged and then disappeared within 3 minutes. Representative stills from live imaging of mCherry-PCNA and EGFP-Rpb3 (a subunit of RNAPII) in control or Cdc7i-injected embryos during S phase 13. Recombinant mCherry-PCNA was co-injected with control or Cdc7i. Time relative to the start of S phase 14 is indicated in minute:second. Scale bars, 5 μm. All images are maximal projections.

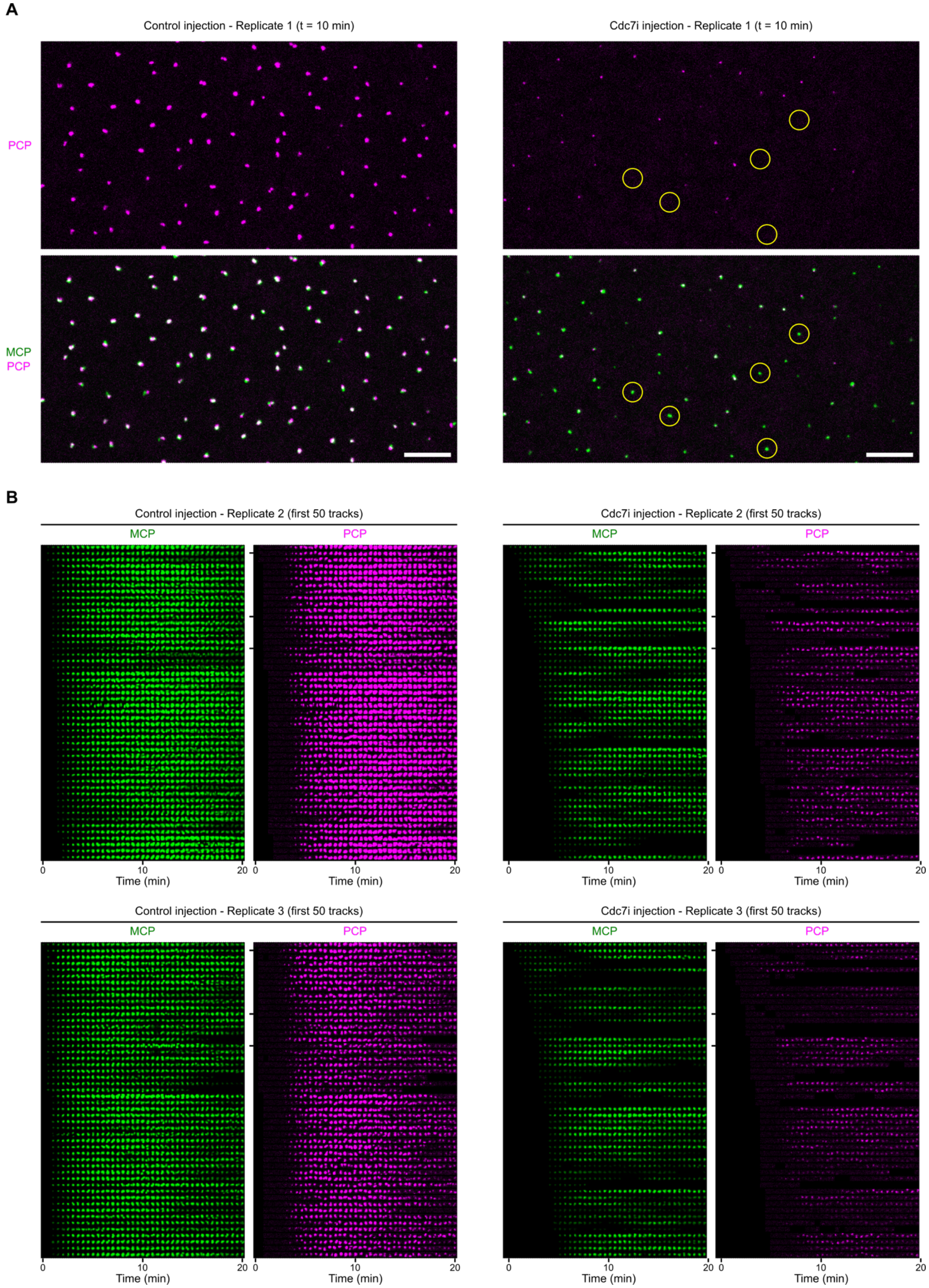


**Supplementary Figure 5. Additional data related to Figure 5. A** Snapshots from live imaging of MCP-GFP and mCherry-PCP in control or Cdc7i-injected embryos with the *hbP2-24xMS2-lacZ-24xPP7* reporter. Frames at 10 minutes after the initial apperance of MCP foci in the same embryo are shown. Yellow circles highlight the MCP foci that lack associated PCP foci. The embryos shown here are the same biological replicates as those in Figure 5B. Scale bars, 10 um. **B** Montages displaying signals from the transcriptional reporter in additional biological replicates of control or Cdc7i-injected embryos during S phase 14. The top 50 spots with the earliest MCP appearance are displayed.

| **Supplementary Table 1.** ***Drosophila melanogaster*** **lines used in this study** | |
|---|---|
| **Genotype** | **Source** |
| *w, sfGFP-Brd4, mCherry-Rpb1* | Cho and O'Farrell, 2023 |
| *w, mKate2-Brd4* | This paper |
| *w, Cdc7-EGFP* | Seller and O'Farrell, 2018 |
| *w, mKate2-Brd4, Cdc7-EGFP* | This paper |
| *w, mNeonGreen-CBP* | This paper |
| *w, mKate2-Brd4, Cdc7-EGFP;; UASp-shRNA.w* | This paper; Bloomington Drosophila Stock Center (#35573) |
| *w, mKate2-Brd4, Cdc7-EGFP;; UASp-shRNA.nej* | This paper; Bloomington Drosophila Stock Center (#36682) |
| *w, mKate2-Brd4, Cdc7-EGFP;; Mat-tub-Gal4* | This paper; Bloomington Drosophila Stock Center (#7063) |
| *w; EGFP-Rpb3; nos-MCP-mCherry* | Cho et al., 2022 |
| *yw; hbP2-MS2-lacZ* | Bloomington Drosophila Stock Center (#60338) |
| *w; nos-NLS-mCherry-PCP, His2Av-eBFP2/CyO; MCP-GFP* | Fukaya et al., 2017 |
| *hbP2-MS2-lacZ-PP7* | Fukaya et al., 2017 |
